# A Chromosome-scale draft genome sequence of horsegram (*Macrotyloma uniflorum*)

**DOI:** 10.1101/2021.01.18.427074

**Authors:** Kenta Shirasawa, Rakesh Chahota, Hideki Hirakawa, Soichiro Nagano, Hideki Nagasaki, Tilak Sharma, Sachiko Isobe

**Affiliations:** Kazusa DNA Research Institute, 2-6-7 Kazusa-kamatari, Kisarazu, Chiba, 292-0818, Japan; Department of Agricultural Biotechnology, CSK Himachal Pradesh Agricultural University, Palampur, HP, 176062, India; ICAR–Indian Institute of Agricultural Biotechnology, Ranchi, Jharkhand, 834010, India; Forest Tree Breeding Center, Forestry and Forest Products Research Institute, 3809-1 Ishi, Juo, Hitachi, Ibaraki, 319-1301, Japan

**Keywords:** Horsegram, *Macrotyloma uniflorum*, Draft genome sequencing

## Abstract

Horsegram [*Macrotyloma uniflorum* (Lam.) Verdc.] is an underutilized warm season diploid legume (2n=20, 22), It is consumed as a food legume in India, and animal feed and fodder in Africa and Australia. Because of its ability to grow under water-deficient and marginal soil conditions, horsegram is a preferred choice in the era of the global climatic change. In recognition of its potential as a crop species, we generated and analyzed a draft genome sequence for HPK-4. The genome sequences of HPK-4 were generated by Illumina platform. Ten chromosome-scale pseudomolecules were created by aligning scaffold sequences onto a linkage map. The total length of the ten pseudomolecules were 259.2 Mb, covering 89% of the total length of the assembled sequences. A total of 36,105 genes were predicted on the assembled sequences, and 14,736 were considered to be horsegram specific genes by comparative analysis with *Phaseolus. vulgaris*, *Vigna. angularis*, *Lotus. japonicus* and *Arabidopsis. thaliana*. The results of macrosynteny analysis suggested that the genome structure of *V. angularis* is more similar to horsegram than that of *P. vulgaris.* Diversity analysis in the 91 accessions of horsegram with dd-RAD-Seq analysis indicated narrow genetic diversity among the horsegram accessions. This is the first attempt to generate a draft genome sequence in horsegram and will provide a reference for sequence-based analysis of the horsegram germplasm to elucidate the genetic basis of important traits.

## Introduction

Horsegram [*Macrotyloma uniflorum* (Lam.) Verdc.], an underutilized warm season diploid legume (2n=20, 22) belonging to tribe Phaseoleae, family Fabaceae is mainly cultivated in semi-arid regions of the world. On the Indian subcontinent, horsegram is primarily consumed as a food legume, whereas in Africa and Australia it is mainly grown for use as a concentrated animal feed and fodder. Horsegram is a self-pollinated plant species thought to originate in Africa, because most of its 32 wild species exist there (Verdcourt, 1971), and the North Western Himalayan region is considered its secondary center of origin (Arora and Chandel, 1972). Horsegram may have been domesticated as a *M*. *uniflorum* var. uniflorum in the Southern part of India, but its probable progenitor *M. axillare* has not been reported in India. Therefore, the process by which cultivated horsegram was domesticated from its wild ancestors has not yet been established (Chahota et al. 2013).

Because of its ability to grow under water-deficient and marginal soil conditions, horsegram is a preferred choice in the era of global climate change. As a result of its medicinal importance and its ability to thrive under drought-like conditions, the US National Academy of Sciences has identified this legume as a potential food source for the future (National Academy of Sciences, 1978). Horsegram contains 16%–30.4% protein (Patel et al., 1995), which constitutes an important source of dietary protein for the largeundernourished population in south Asia. In addition, the seeds are a rich source of lysine and vitamins (Goplan et al., 1989), and its antioxidant, antimicrobial and unique antilithiatic properties make it a food of nutraceutical importance (Reddy et al., 2005; Manisha et al., 2020a, 2020b).

The existence of many wild and unsolicited characteristics makes horsegram a less favorable legume for commercial cultivation, although it does possess numerous attributes that make it a potential food legume for warm arid regions. In addition, this crop species has not benefited much in terms of development of genetic and molecular tools for its genetic enhancement. Despite these limitations, the crop has potential to provide nutritional food security to the resource poor farmers of the developing world during climactic changes and regional population surges. In recognition of the potential of this food legume species, we generated and analyzed a draft genome sequence for HPK-4, a horsegram cultivar commercially released by CSK Himachal Pradesh Agricultural University (HPAU), Palampur, India. This variety, which has a dark brown seed color, is under cultivation in many parts of the Northwestern Indian Himalayan region. It is resistant to anthracnose (*Colletotrichum truncatum*) and tolerant to abiotic stresses such as drought, salinity, and heavy metals. This is the first attempt to generate a draft genome sequence of this “orphan”, but it is an important food legume species and will provide a reference for sequence-based analysis of the horsegram germplasm to elucidate the genetic basis of important traits.

## Results

### Whole genome sequencing and assembly of horsegram

The genome sequences of the horsegram variety, HPK-4, were generated from a paired end (PE) library by Illumina HiSeq 2000 with a total length of 37.9 Gb (Table S1). Using Jellyfish program (Marçais and Kingsford, 2011), the genome size of HPK-4 was estimated as approximately 343.6 Mb (Fig. S1).

The Illumina PE reads were assembled by SOAPdenovo2 (Luo et al., 2012) with kmer = 61 or 81, and contigs were generated with a total length of 352.2 Mb (kmer = 81) and 389.3 Mb (kmer = 61 M) (Table S2). The contigs constructed with kmer = 81 were selected and scaffolded with mate pair (MP) reads by using SSPACE2.0 (Boetzer et al., 2011). The number of generated scaffolds was 6,227 after gap filling by GapFiller (Boetzer and Pirovano, 2012) with the PE reads and excluding contamination. The total length of scaffolds was 297.1 Mb (Table S2, Assembly 1), which was approximately 55– 92 Mb shorter in length than the estimated genome size of HPK-4.

We speculated that the observed shorter length of the total scaffolds may have been due to miss-integration of repeat sequences by SSPACE, and we therefore performed the subsequent assemblies with PE and MP reads using two programs, Platanus (Kajitani et al., 2014) and MaSuRCA (Zimin et al., 2013). The total ATGC lengths of the scaffolds in the three assemblies were not significantly different in the three assemblies, 284.4 Mb in SOAPdenovo-SSPACE (Assembly 1, before excluding contamination; Table S2), 277.0 Mb in Platanus (Assembly 2), and 299.1 Mb in MaSuRCA (Assembly 3). The Assembly 1 was used for subsequent analysis, and gaps were closed with Illumina synthetic long-reads (SLRs) by GM closer (Kosugi et al., 2015). The total length of the resultant assembly (Assembly 5) was 295.7 Mb.

The results of benchmarking universal single-copy orthologs (BUSCOs) analysis (Simão et al., 2015) identified 93.1% of the complete BUSCOs (data not shown) in the Assembly 5. We therefore considered that Assembly 5 covered most of the coding regions of the horsegram genome. The sequences having a length of less than 500 bp were excluded from Assembly 5, and the remaining sequences were designated as MUN_r1.1.

### Linkage map and pseudomolecule construction

To construct chromosome-scale genome sequences, a single nucleotide polymorphism (SNP) linkage map was created with the 214 F_2_ progenies. The female parent of the F_2_ progenies was HPK-4. The male parent was initially considered to be HPKM-193, but this assignment was later found to be wrong when the whole genome sequences of HPK-4, HPKM-193 and eight F_2_ progenies were compared. The candidate SNPs segregating in the F_2_ progenies were identified by mapping the whole genome Illumina sequences of the eight randomly selected F_2_ progenies (Table S1) onto MUN_r1.1. A total of 2,942 SNPs were identified, and 1,378 SNPs were successfully genotyped by target amplicon sequencing (TAS) analysis in 214 F_2_ progenies. Of these, 1,263 SNPs were mapped onto the ten linkage groups with a total length of 980 cM (Table S3). A total of 219 scaffolds in MUN_r1.1 were then aligned onto the linkage map (Fig.1; Table S3; Table S4).

**Figure 1.**
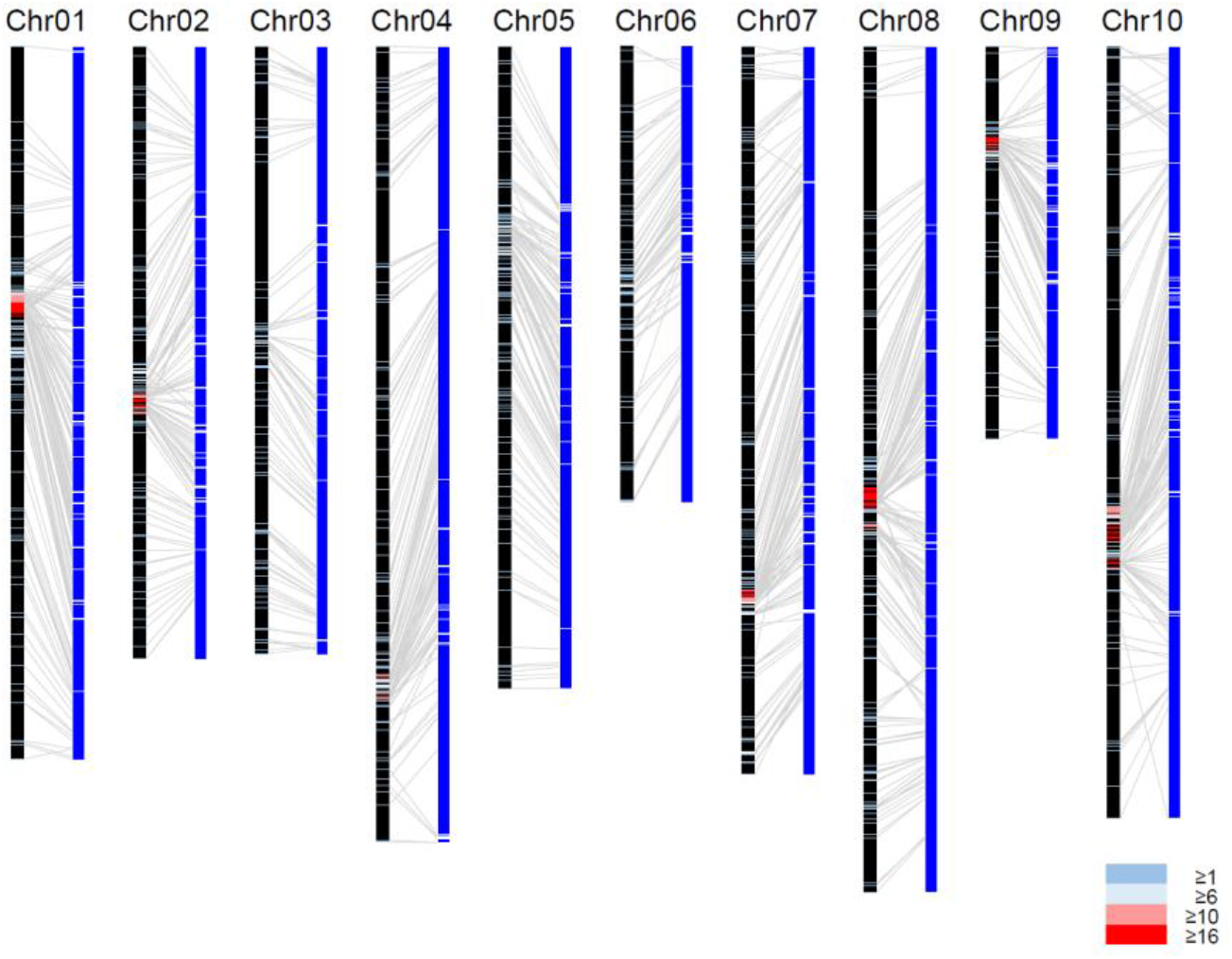
Anchoring the horsegram genome assembly to the genetic linkage map. The linkage groups (left black bars) and 219 anchored MUN_r1.1 scaffolds (right blue bars) with 1,263 SNPs. The crossbars on the linkage groups show the positions of mapped SNPs. Blue, aqua, pink and red colors represent numbers of mapped SNPs per cM of 1~5, 6~10, 10~15 and ≧16, respectively.

During the process of alignment, two scaffolds were discovered to be miss-scaffoldings and split. The revised set of the scaffolds was designated as MUN_r1.11 (Table 1 and Table S1). The number of sequences of MUN_r1.11 was 3,495, with a total length of 294.7 Mb and N50 length of 2.8 Mb. The aligned scaffolds on the linkage map were connected with 10,000 Ns for the construction of chromosome-scale pseudomolecules. The total length of the 10 pseudomolecules was 259.2 Mb, with an N50 length of 28.2 Mb (Table 1). The length of pseudomolecules ranged from 15.5 Mb (chr09) to 33.4 Mb (chr08, Table S5). When the total length of the A, G, T and C bases was compared, the 10 pseudomolecules covered 89% of the scaffolds in MUN_r1.11. The ratios of complete BUSCOs identified in MUN_r1.11 and the 10 scaffolds were 93.1% and 87.4%, respectively. Most of the complete BUSCOs were identified as single copies, suggesting that the rate of duplication in the coding regions of the assembled genomes was low.

**Table 1.** Statistics of the Horsegram genome assembly and CDS.

|  | MUN_r1.11 | MUN_r1.11 | MUN_r1.1_cds |
| --- | --- | --- | --- |
|  | Genome / Scaffolds | Genome / Pseudomolecules | CDS |
| Number of Sequences | 3,497 | 10 | 36,105 |
| Total length (bp) | 294,688,765 | 259,245,825 | 38,820,013 |
| Average length (bp) | 84,269 | 25,924,583 | 1,075 |
| Maximum length (bp) | 9,844,273 | 33,386,276 | 15,732 |
| Minimum length (bp) | 500 | 15,505,026 | 150 |
| N50 length (bp) | 2,818,555 | 28,154,654 | 1,488 |
| Total length of AGTC (bp) | 286,841,254 | 255,003,950 |  |
| Gaps (bp) | 7,847,511 | 4,241,875 | - |
| GC% | 30.8 | 30.5 | 43.8 |
| Repeat % | 28.99497136 | - | - |
| BUSCOs |  |  |  |
| Complete | 1340 (93.1%) | 1259 (87.4%) | 1313 (91.2%) |
| Complete Single-Copy | 1252 (86.9%) | 1181 (82%) | 1208 (83.9%) |
| Complete Duplicated | 88 (6.1%) | 78 (5.4%) | 105 (7.3%) |
| Fragmented | 26 (1.8%) | 31 (2.2%) | 23 (1.6%) |
| Missing | 74 (5.1%) | 150 (10.4%) | 104 (7.2%) |
| Number of complete genes | - | - | 35,508 |
| Number of partial genes | - | - | 597 |

### Repetitive sequences

Repetitive sequences in the genomes were identified by RepeatMasker 4.0.7 for known repetitive sequences registered in Repbase and *de novo* repetitive sequences defined by RepeatModeler 1.0.11. A total of 50.2 Mb of repetitive sequences were identified on the assembled genome, occupying 29% of the total length (Table S6). The ratio of repetitive sequences varied among the chromosomes, ranging from 22.5% (chr06) to 36.8% (chr01). Of the identified repetitive sequences, the sequences registered in Repbase were found on 12.0% of the assembled genome, whereas unique repetitive sequences, i.e., not registered in Repbase, were located on 17.0% of the assembled genome. A total of 74,362 simple sequence repeats (SSRs) were identified in MUN_r1.11 with an average frequency of 0.21 SSR per 100 kb (Table S7). The highest SSR frequency of 0.66 SSR per 100 kb was observed in chr06, and was almost three times higher than that of chr03 and chr08 (0.22 SSR per 100kb).

### Transcript sequencing, gene prediction and annotation

A total of 485 M transcript Illumina reads were obtained from seedlings, leaves, roots, flowers and young pods of HPK-4 (Table S1; Fig. S2). Gene prediction was performed for MUN_r1.11 by BRAKER1 v1.9 (Hoff et al., 2016) with the obtained transcript and published transcript sequences by Bhardwaj et al. (2013). A total of 46,095 gene sequences were predicted on the assembled genome with a total length of 48.3 Mb (Table S8a). After removal of transposon elements (TEs) and pseudo and short gene sequences, the number of gene sequences was 36,105. These complete and partial gene sequences were designated as MUN_r1.1_cds (Table 1). The ratio of complete BUSCOs identified on MUN_r1.1_cds was 91.2%. Of the 36,105 gene sequences, 35,508 were classified as complete genes, and 597 were classified as partial genes. The coding sequences (CDSs) were further tagged with “f” (full similarity), “p” (partial similarity) and “d” (domain) according to the similarity level against the NR database (f: E-values ≤1E-20 and identity ≥70%; p: E-values ≤1E-20 and identity <70%) and InterPro database (d: E-values ≤1.0; Table S8b). Of the 36,105 complete and partial genes, 21,471 (59.4%) sequences were tagged with “f” and 6,692 (18.5%) were tagged with “p”. The number of predicted gene sequences tagged with “d” was 24,575 (68.1%).

The total number of putative tRNA genes in the assembled genomes (MUN_r1.11) was 690, almost the same as the numbers for *Phaseolus vulgaris* (681), *Vigna angularis* (667) and *A. thaliana* (699, Table S9). The total number of putative rRNA genes identified in the genome was 139, which was again the same as the number in the *P. vulgaris* genome.

### Comparative and phylogenetic analysis with other plant species

The predicted gene sequences in MUN_r1.1_cds were clustered with other plant species (*P. vulgaris*, *V. angularis*, *L. japonicus* and *A. thaliana*) for comparison at the protein sequence level. A total of 73,457 clusters were generated using the program CD-HIT (Fu et al., 2012, Table S10). Of the 36,105 putative gene sequences, 21,369 (59.2%) genes were clustered with other plant species, and 14,736 (40.8%) were considered to be horsegram-specific genes (Fig. 2). A total of 3,738 (10.4%) horsegram gene sequences were clustered with 3,864 *P. vulgaris* and 3,713 *V. angularis* genes, which were considered to be Millettioids-specific genes. Common genes in legumes were identified for 6,550 (18.1%) horsegram gene sequences, based on clusters with *P. vulgaris*, *V. angularis* and *L. japonicus*.

**Figure 2.**
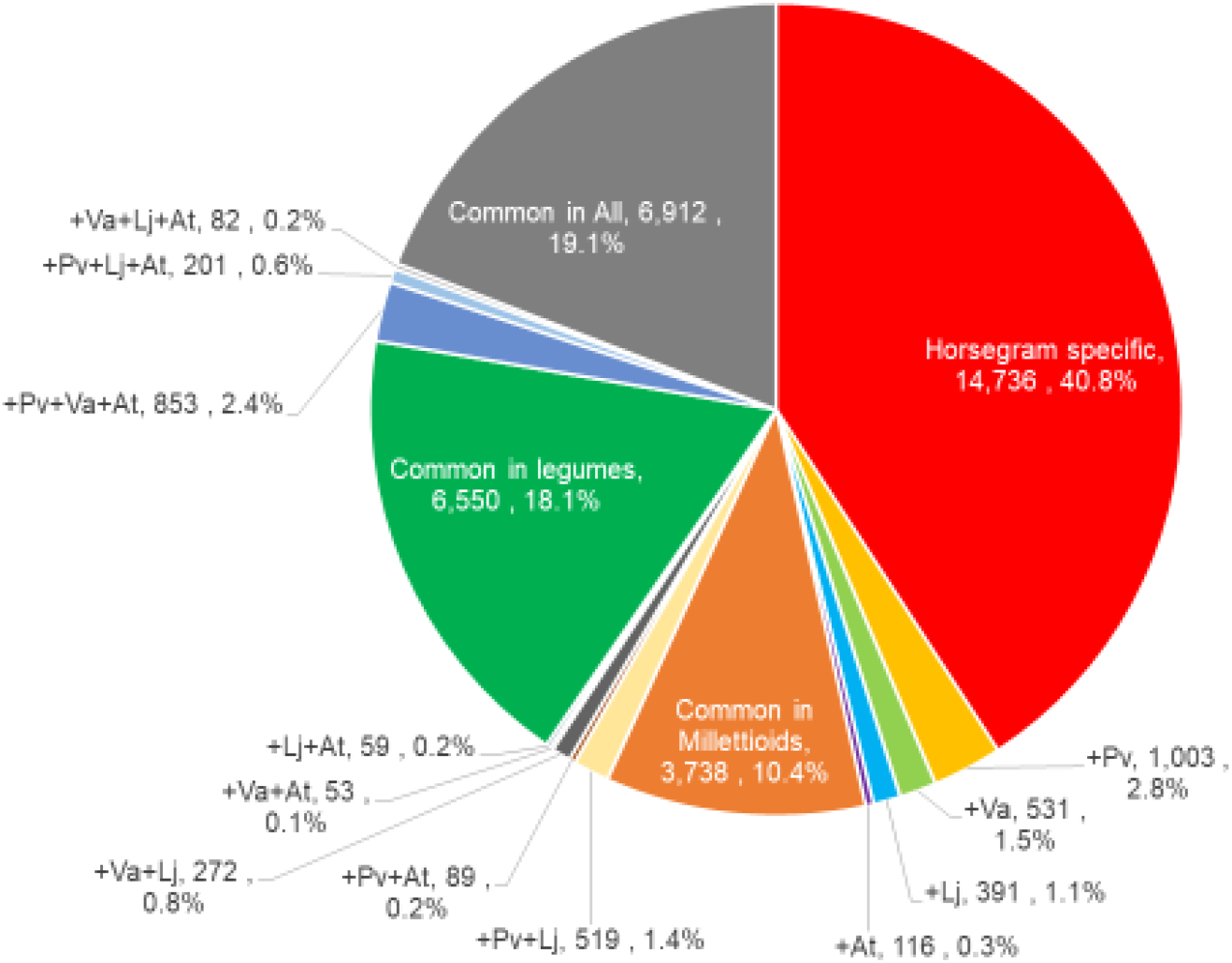
Ratios of genes of horsegram (MUN_r1.1_cds) clustered with four other plant species. Pv, Va, Li and At represent genes of *P. vulgaris* (Pvulgaris_218_v1.0), *V. angularis* (Vangularis_vl.al), *L. japonicus* (Lj_r3.0), and *A. thaliana* (Araport11), respectively.

Functional analysis was performed for MUN_r1.1_cds by classifying 36,105 putative genes into the gene ontology (GO) and “euKaryotic clusters of Orthologous Groups” (KOG) databases. A total of 24,699 (68.4%) putative genes were annotated with GO categories including 9,086 (25.2%) genes involved in biological processes, 4,127 (11.4%) genes coding for cellular components and 1,377 (38.7%) genes associated with molecular functions (Fig. S3). The ratio of annotated genes was less compared to other species. The species with a ratio of classified GO categories most similar to that of horsegram was *L. japonicus*. A total of 18,630 (51.6%) putative genes showed significant similarity to genes in the KOG database (Fig. S4). As in the results for GO, the ratio of hit genes was less than for the other four species.

### Genes related to drought tolerance

The candidates for genes related to drought tolerance were searched against the 199 genes registered in the Drought Stress Gene Database (DroughtDB; Alter et al. 2014). A total of 158 horsegram genes showed significant similarity to the 78 genes in the DroughtDB (Table S11). The most frequently hit gene was *ABCG40*, which functions as an ABC-transporter and showed significant similarity to the 14 horsegram genes. *OST1/SRK2E* and *AtrbohF* were also frequently identified, with hits to 7 and 6 horsegram genes, respectively. Of the 158 genes, 93 showed the same domain sequences as the *A. thaliana* gene, and 52 were similar to the genes registered in the Plant Stress Gene Database (PSGD: http://ccbb.jnu.ac.in/stressgenes/frontpage.html). These genes were indicated to have a greater likelihood of being candidate genes related to drought tolerance.

### Diversity analysis in genetic resources

Genetic diversity in 91 cultivated horsegram accessions and one *M. axellare* accession, which is a wild relative of horsegram, being maintained at CSK-HPAU were investigated based on dd-RAD-Seq analysis (Table S1). The two accessions, IC139449 and IC547543, were excluded from the further analysis because of the small number of obtained reads. The reads were mapped on the assembled genome (MUN_r1.11). The mapped ratio of the reads onto the genome ranged from 80% to 90% in most of the accessions (Fig. S5). However, *M. axellare* and one horsegram accession (IC313367) showed low mapping ratios of 17% and 55%, respectively. *M. axellare* was excluded from further analysis because of its low mapping ratio.

A total of 277 SNPs were identified in the remaining 89 accessions across the genome (Table S12). The NJ tree revealed that the 89 accessions were classified into two clusters (Fig. 3). Cluster 1 included varieties bred in the CSK-HPAU, which are prefixed with “HPK”. Most of the HPK varieties showed very close genetic relations and formed a subcluster (HPK Cluster); the single exception was HPK-4.

**Figure 3.**
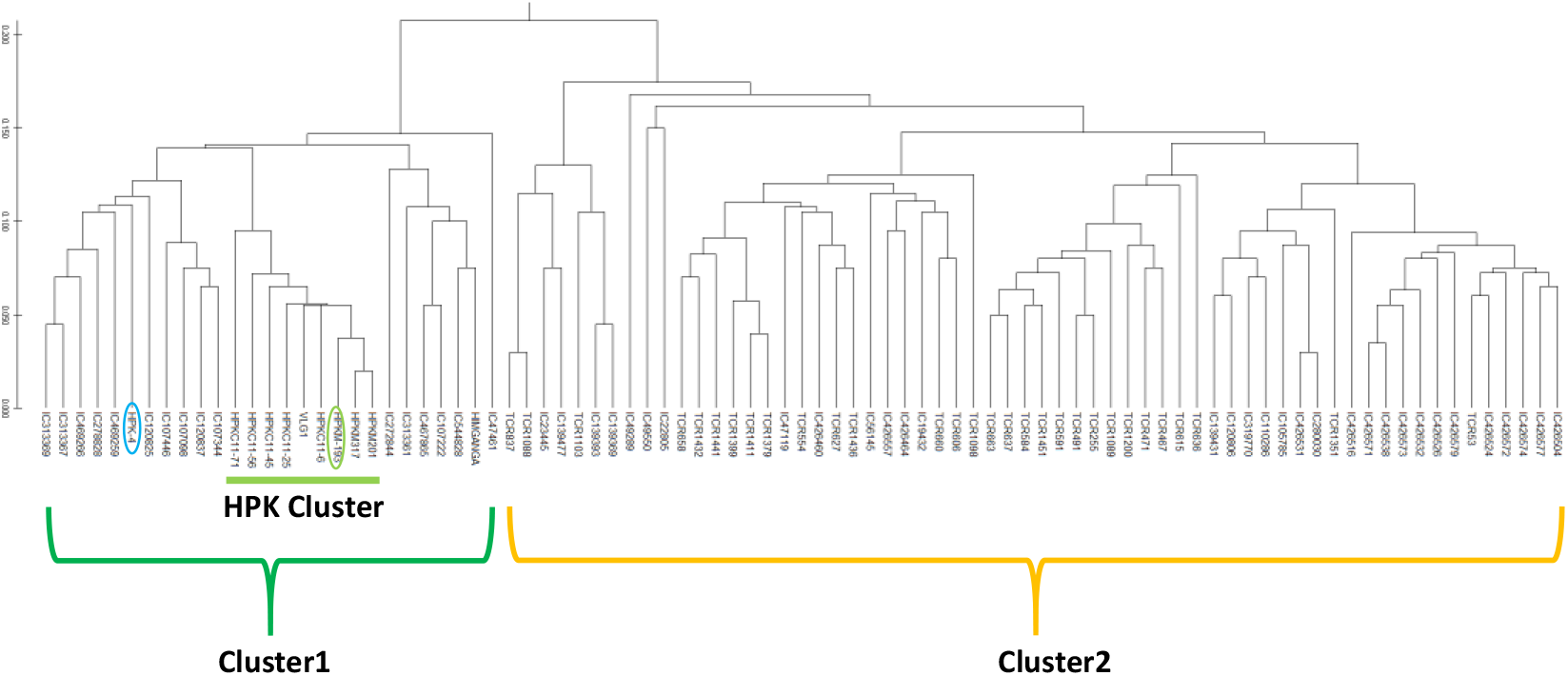
A phylogenetic tree of the 89 horsegram accessions based on 277 SNPs. HPK-4, used in the reference genome construction; HPKM-193, the obtained whole genome sequences are circled with blue and green lines.

### Whole genome structure in horsegram

The distributions of repetitive sequences in the horsegram genome are shown in Fig. 4A. The repetitive sequences were frequently observed in the midsection of each chromosome, and the tendency was more pronounced in horsegram-specific sequences. The ratio of repetitive sequences commonly observed in all five species was quite low, suggesting the uniqueness of repetitive sequences compared to the gene sequences. The gene sequences commonly observed between horsegram and other compared species tended to distribute in the two end regions of the chromosomes (Fig. 4B). On the other hand, horsegram-specific gene sequences were distributed more uniformly across the genome, suggesting the unique structure of the horsegram genome.

**Figure 4.**
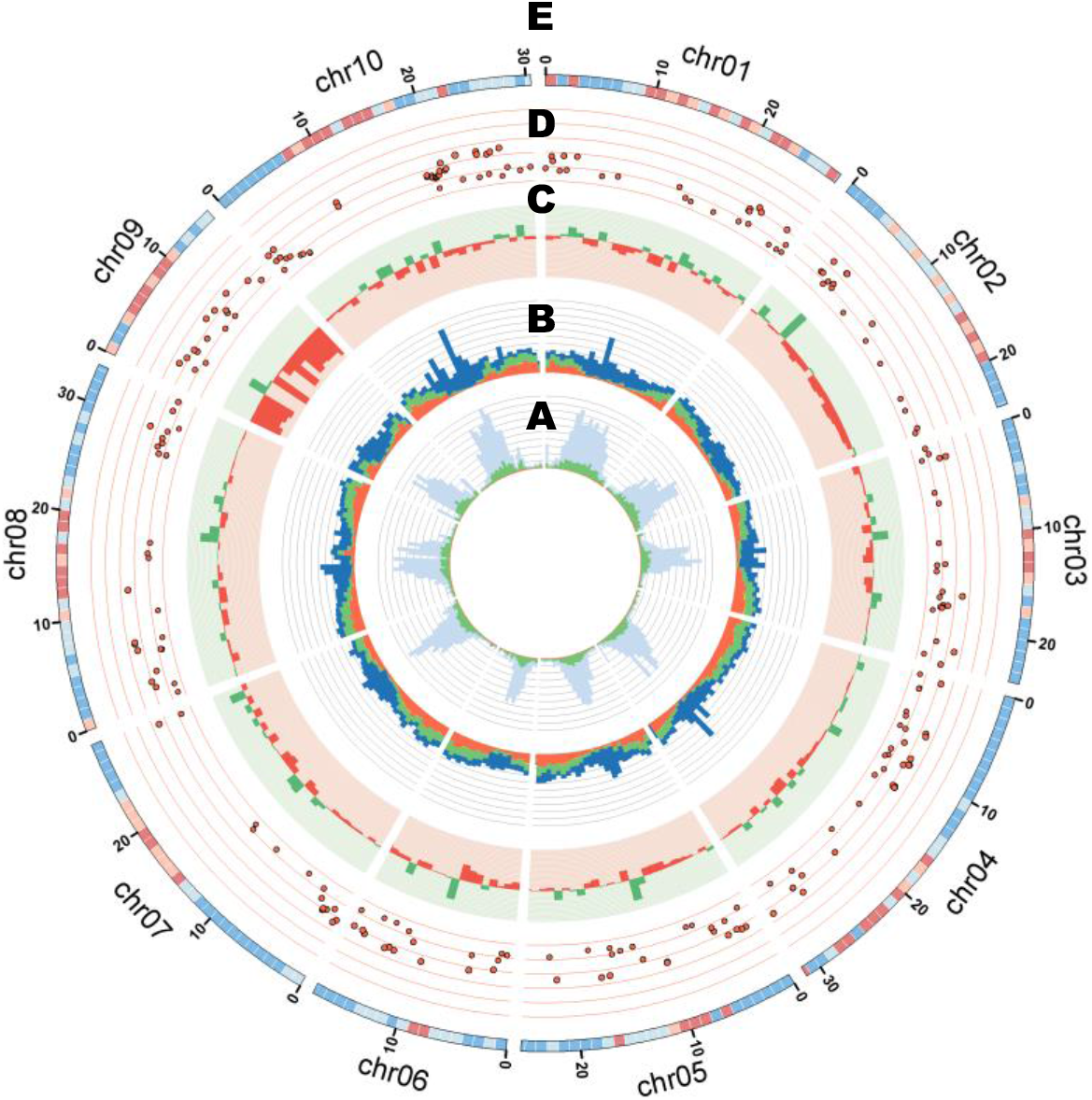
Graphical view of the horsegram genome structure. **A:** Ratios of repetitive sequences in 1 Mb windows. Blue bars represent horsegram-specific sequences. Green bars show sequences commonly observed in horsegram and three other legume species*, P. vulgaris*, *V angularis* and *L. japonicus*. Red bars show sequences commonly observed in the four legume species and *A. thaliana*. **B:** Numbers of the predicted horsegram genes (MUN_r1.1_cds) in 1 Mb windows. The bar colors are the same as in (A). **C:** CNV distribution in a 1 Mb window. Green and red dots show log2 ratio plus and minus values, respectively. **D:** Pi value and positions of the 255 SNPs identified in the 89 horsegram accessions by using dd-RAD-Seq. The distance between horizontal lines represents a Pi value of 0.1. **E:** SNP density identified among the F2 mapping population based on whole genome re-sequencing of the eight F2 progenies. Blue, pale blue, pale pink and pink indicate numbers of SNPs less ≦50, ≦100, ≦150 and 150< in 1 Mb windows, respectively.

Copy number variations (CNVs) of one horsegram accession, HPKM-193, were detected against the HPK-4 genome (Fig. 4C) based on the whole genome sequence reads of HPKM-193. CNVs with a minus log2 ratio were particularly observed on chr09 and chr02. SNP density mapped on the linkage map is illustrated in Fig. 4E. As in the case of the CNVs, a distribution bias was observed in the SNPs of HPKM-193; however, this bias was not similar to that in CNVs. A higher SNP density was observed in the midsection in most of the chromosomes. Chr06 showed less variation than the other chromosomes.

Of the 277 SNPs identified among 89 horsegram accessions, 255 were located across genome sequences of 10 chromosomes (Fig. 4D). In each chromosome, the SNPs tended to be identified in the regions where common putative genes of horsegram and the other compared species were located, particularly for chr04, chr07, chr08 and chr10. It was considered that the differing trends in variable distribution reflected the presence of varying degrees of selection pressure in the horsegram germplasm resources in Himachal Pradesh.

### Comparative and phylogenetic analysis with other legume species

The genome structure of horsegram was compared with those of *P. vulgaris*, *V. angularis* and *L. japonicus* by aligning homologous sequence pairs along each pseudomolecule (Fig. 5a). The comparison revealed obvious syntenic relations between horsegram and the other legume species. Clear relations were observed with warm season legume, *V. angularis*, one on one relationships were observed between horsegram chr02 (Mun_chr02) and *V. angularis* chr09 (Va_chr09), Mun_chr04 and Va_chr02, Mun_chr06 and Va_chr10, Mun_chr07 and Va_chr08, Mun_chr08 and Va_chr04, and Mun_chr09 and Va_chr05. The syntenic relations with *P. vulgaris* were slightly more complex than those with *V. angularis,* and those with cool season legume *L. japonicus* were more fragmented.

**Figure 5:**
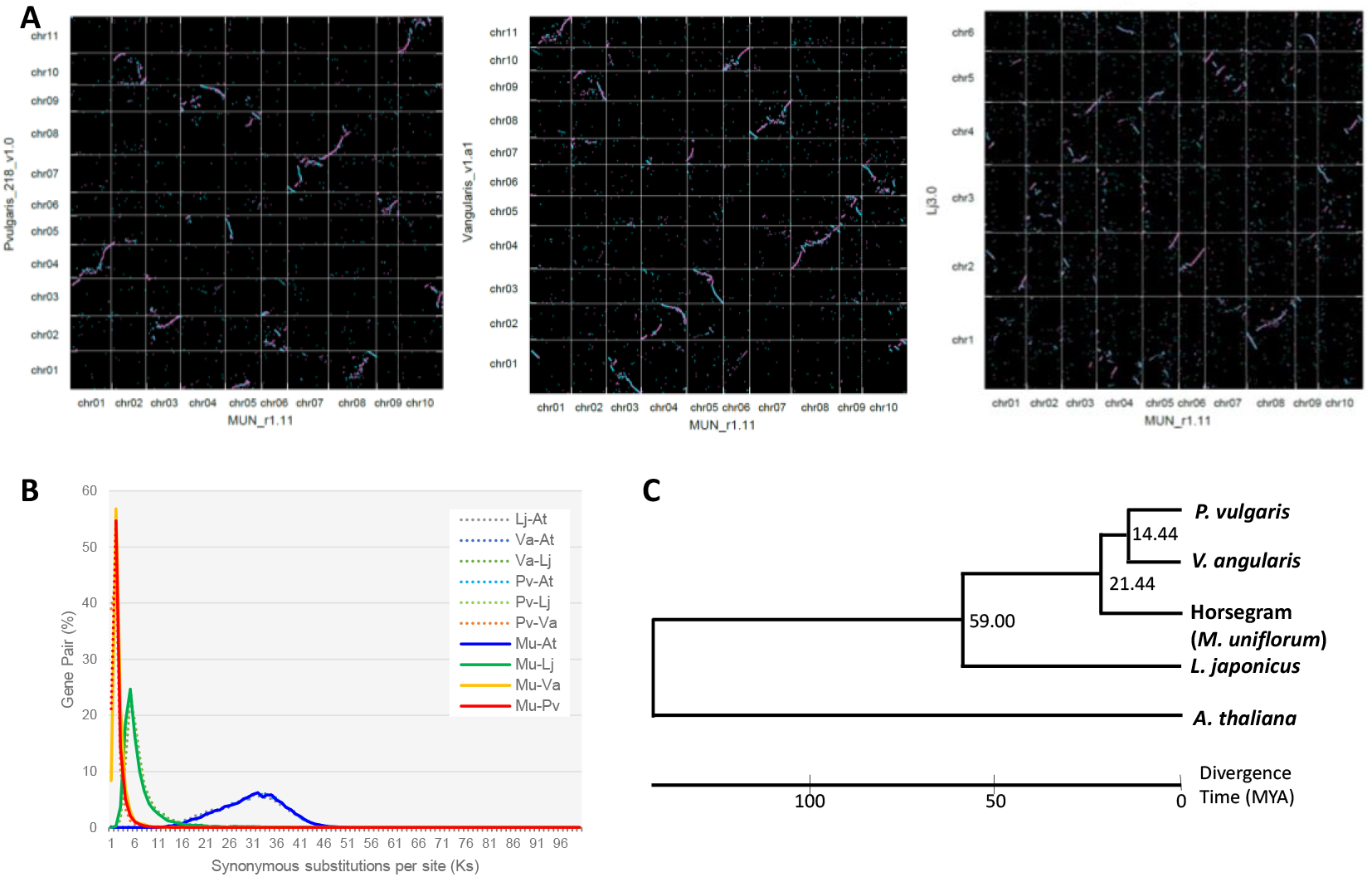
Comparative and phylogenetic analysis with other legume species. **A:** Graphical view of syntenic relationships between horsegram *and P. vulgaris* (left), *V. anagularis* (middle) *and L. japonicus* (right). Pink and blue dots show homologous sequences of MUN_r1.11 with forward and reverse directions against the reference sequences. **B:** Distribution of Ks values of orthologous gene pairs in horsegram (Mu) and the four plant species, *P. vulgaris* (Pv), *V. anagularis* (Va), *L. japonicus* (Lj), and *A. thaliana* (At). **C:** A phylogenetic tree of 4,154 common single copy genes of the four legume species and *A. thaliana*.

Synonymous substitutions per site (Ks) were estimated by comparison of gene pairs in each combination of horsegram, *P. vulgaris*, *V. angularis*, *L. japonicus* and *A. thaliana* (Fig. 5b). The similar distributions of horsegram, *P. vulgaris* and *V. angularis* indicated the close relations among the three species. The ratio of gene pairs showing Ks values less than 0.1% was 21.1% between horsegram and *P. vulgaris*, and 8.4% between horsegram and *V. angularis*, suggesting that there was a closer relationship between horsegram and *P. vulgaris* at the gene level.

A neighbor-joining (NJ) tree was created with 4,154 single copy orthologous genes identified in the four legume species. *A. thaliana* was used for the outgroup (Fig. 5c). Among the three legume species in Millettioid, *P. vulgaris* and *V. angularis* shared closer relation than horsegram. When the divergence time between *L. japonicus* and *P. vulgaris* was considered to be 59 million years ago (MYA) (TimeTree; http://www.timetree.org/), it was estimated that horsegram diverged from *P. vulgaris* and *V. angularis* some 21.44 MYA. The divergence time between *P. vulgaris* and *V. angularis* was estimated as 14.44 MYA.

## Discussion

In the present study, we reported *de novo* whole genome assembly in horsegram. This is the first report of de novo whole genome assembly in the genus *Macrotyloma*. The genome size of horsegram was estimated to be 343.6 Mb, but the total length of the assembled genome was 294.7Mb, suggesting that approximately 48.9 Mb of sequences were missing from the assembly. Meanwhile, 96.1% complete BUSCOs were identified in the assembled genome of MUN_r1.11, including 86.9% of single copies and 6.1% of duplicated copies. Therefore, we assumed that most of the genic regions were not missing in the assembly, and the missing 48.9 Mb of sequences might be repetitive sequences. In this study, we used Illumina sequencing platforms for genome assembly. Adding long reads sequences, such as PacBio or Nanopore reads, might help for future improvement of the assembly. Nevertheless, the genome sequences were assembled on the chromosome scale, and undoubtedly the created sequences will contribute to the advancement of genetic and genomic studies in horsegram and the genus *Macrotyloma*.

*De novo* assembly in transcript sequences from shoot and root tissues was previously reported by Bhardwaj et al. (2013). In their report, 26,045 assembled transcripts showed significant BLAST hits against other species, while 3,558 transcripts showed no hits. In our study, 36,105 protein coding genes were predicted on the assembled genome, and 14,736 (40.8%) were assumed to be horsegram-specific genes by clustering analysis with the three legume species and *A. thaliana*. We identified more comprehensive gene sequences based on assembled genome, especially in horsegram-specific gene sequences.

Horsegram is considered one of the most drought-tolerant legume crop species. Personal investigation showed that plants can survive for more than 20 days without watering under controlled conditions. A study by Bhardwaj et al. (2013) described a transcriptome analysis conducted for eight shoot and root tissues of a drought sensitive (M-191) and a tolerant (M-249) genotype of horsegram under controlled and drought stress conditions and identified some important genic regions responsible for drought tolerance. In the present study, we listed 158 candidate drought tolerance genes by BLAST search against the 199 genes registered in the DroughtDB. Although further investigation is of course required, it is expected that the assembled genomes in this study will contribute to a deeper understanding of drought tolerance in horsegram.

Only a few species, horsegram and *M. geocarpum* are used as crops in in gen*us Macrotyloma*. It was speculated that the domestication of horsegram occurred in India twice, in northwestern India in 4000 BP and the Indian Peninsula in 3500 BP (Fuller and Murphy, 2018). In addition, horsegram is known to possess narrow genetic diversity as revealed in molecular analysis (Sharma et al., 2015). In this study, we investigated the genetic diversity in 89 horsegram accessions by using dd-RAD-Seq analysis. The number of identified SNPs was only 277. In addition, the phylogenetic analysis revealed that the accessions collected and/or bred in Himachal Pradesh fell in one cluster (HPK cluster in Fig. 3). Although fewer variants were identified, the morphological diversity was observed among the varieties in the HPK cluster (Chahota et al. 2005). It was also an interesting observation that there were a larger number of CNVs of HPK-193 against the HPK-4 genome in chr09. Further investigation on the association between the variants and morphological traits would identify causal variants for agriculturally important morphological traits.

Horsegram belongs to the subtribe Phaseolinae in the Milletttioids clade along with *P. vulgaris* and *V. angularis*. Based on phylogenetic comparison of 4,154 single orthologous gene sequences, the divergence time between horsegram and *P. vulgaris* / *V. angularis* was estimated as 21. 44 MYA. The genetic distance between horsegram and either *P. vulgaris* or *V. angularis* is greater than the genetic distance between *P. vulgaris* and *V. angularis* themselves, and the results are in consonance with a previous study based on a comparison of eight chloroplast regions (Stefanović et al., 2009). However, the estimated divergence time of 21.44 MYA in this study was older than that of 7.0~10.5 MYA reported by Stefanović et al.

In this study, we have provided a first draft genome assembly of horsegram cultivar (HPK-4) and investigated feature of the horsegram genome and gene sequences as well as genetic diversity in the accessions. It will help in the establishment of an efficient breeding program for horsegram by integrating conventional breeding with marker-based biotechnological tools. Finally, the genomic information revealed in this study can be applied to the improvement of other disadvantageous food legumes.

## Experimental procedures

### Whole genome sequencing and assembly

A horsegram variety, HPK-4, bred at CSK HPAU was used for whole genome sequencing. Illumina PE and MP libraries were constructed with expected insert sizes of 500 bp (PE) and 2 Kb, 5 Kb, 10 Kb and 15 Kb (MP). Library sequencing was performed by Illumina HiSeq 2000 and MiSeq (Illumina, San Diego, CA) systems with read lengths of 93 nt or 101 nt (Seq 2000) and 251 nt (MiSeq; Supplementary Table S1). An Illumina SLR library was also constructed with high molecular cellular DNA. The SLR library was constructed by a TruSeq synthetic long-read DNA library prep kit (Illumina), and sequences were generated by Illumina HiSeq 2000 and MiSeq systems with read lengths of 93nt and 251nt, respectively. The SLR reads were synthesized through the TruSPAdes pipeline (Bankevich and Pevzner, 2016).

Three approaches were applied for the PE and MP reads assembly (Supplementary Table S2). The first assembly (Assembly 1) was done using the combination of SOAPdenovo2 (Luo et al., 2012), SSPACE2.0 (Boetzer et al., 2011) and GapFiller (Boetzer and Pirovano, 2012). Scaffolding with the MP reads was performed by using SSPACE2.0, and the gaps of the scaffolds were closed by GapFiller with PE reads. The other two assemblies (Assemblies 2 and 3) were performed with Platanus (Kajitani et al., 2014) and MaSuRCA (Zimin et al., 2013). Gap closing with SLR assembly (Assembly 4) was performed for Assembly 1 by using GMcloser (Kosugi et al., 2015). Potentially contaminated sequences were excluded by using BLASTN searches against the chloroplast and mitochondrial genome sequences of *Arabidopsis thaliana* (accession numbers: NC_000932.1 and NC_001284.2), human genome sequences (hg19, https://genome.ucsc.edu/), fungal and bacterial genome sequences registered in NCBI (http://www.ncbi.nlm.nih.gov), vector sequences in UniVec (http://www.ncbi.nlm.nih.gov/tools/vecscreen/univec/), and PhiX (NC_001422.1) sequences with E-value cutoffs of 1E-10 and length coverage >10%.

The genome size of HPK-4 was estimated by kmer frequency distribution (kmer=17) using Jellyfish ver. 2.1.1 (Marçais and Kingsford, 2011). Assembly quality was assessed with BUSCO sequences by using BUSCO v3.0 (Simão et al., 2015). Repetitive sequences in the assembled genome were identified by RepeatMasker 4.0.7 (http://www.repeatmasker.org/RMDownload.html) for known repetitive sequences registered in Repbase (https://www.girinst.org/repbase/) and *de novo* repetitive sequences defined by RepeatModeler 1.0.11 (http://www.repeatmasker.org/RepeatModele). SSR motifs were identified by MISA mode in SciRoKo software with default parameters (Kofler et al., 2007).

### Linkage map and pseudomolecule construction

An F_2_ mapping population consisting of 214 individuals was used for linkage map construction. SNPs segregating in the F_2_ population were detected by mapping Illumina re-sequence reads of the eight F_2_ individuals onto the assembled genome using Bowtie2 (Langmead and Salzberg, 2012), and calling variants using SAMtools 0.1.19 (Li et al., 2009) and Varscan 2.3 (Koboldt et al., 2012). TAS was performed for genotyping of the identified SNPs according to the methods described in Shirasawa et al. (2016). Illumina MiSeq was used for amplicon sequencing with a read length of 251 nt. Linkage map construction was performed using JoinMap 4 with Kosambi’s mapping function (Van Ooijen, 2006). The assembled genome sequence scaffolds were aligned onto the linkage map for pseudomolecule construction.

### Transcript sequencing, gene prediction and annotation

Total RNA of HPK-4 was extracted from seedlings, leaves, roots, flowers and young pods by using an RNeasy Plant Mini Kit (QIAGEN). RNA libraries were constructed by using a TruSeq standard mRNA HT sample prep kit (Illumina). Library sequencing was performed by an Illumina HiSeq system with a read length of 93 nt. The obtained transcript sequences were assembled by Trinity (Henschel et al., 2012).

Evidence-based gene prediction was performed for the assembled genome sequences by BRAKER1 v1.9 (Hoff et al., 2016) with the obtained and published transcript sequences by Bhardwaj et al. (2013) (SRP029360). *ab initio* gene prediction by Augustus in the MAKER-P pipeline (http://www.yandell-lab.org/software/maker-p.html) was additionally performed with the predicted gene sequences by BRAKER1 and amino acid sequences of *A. thaliana* (Araport11, https://www.arabidopsis.org/), *L. japonicus* (Lr_r3.0; Sato et al., 2008), *P. vulgaris* (common bean, Pvulgaris_218_v1.0) (Schmutz et al., 2014) and *V. angularis* (azuki bean, Vangularis_v1.a1) (Sakai et al., 2015). Evidence-based gene prediction was also performed by mapping the horsegram transcript sequences onto the assembled genome sequences by using the Tophat-Cufflinks pipeline (Trapnell et al., 2009). The results of *ab initio* and evidence-based gene prediction were merged by the MAKER-P pipeline.

TEs were detected by BLASTP searches against the NCBI NR protein database (http://www.ncbi.nlm.nih.gov) with an E-value cutoff of 1E-10. Domain search was performed by InterProScan (Quevillon et al., 2005) against the InterPro database with an E-value cutoff of 1.0 (Mulder et al., 2007) and BLASTP search against the NCBI NR protein database with E-value ≦1E-20. Prediction of transfer RNA genes was conducted using tRNAscan-SE ver. 1.23 with default parameters (Lowe and Chan, 2016). rRNA genes were predicted by BLAST searches (E-value cutoff of 1E-10) with query sequences of *A. thaliana* 5.8S and 25S rRNAs (X52320.1) and 18S rRNA (X16077.1). Functional analysis of CDSs in CSE_r1.1_cds was performed by classifying the sequences into the plant GO slim categories and KOG categories (Tatusov et al., 2003).

### Genes related to drought tolerance

BLASTP search of the 36,105 putative genes was performed against amino acid sequences of *A. thaliana* (Araport11), and hit genes were further used in BLAST searches against the DroughtDB (Alter et al. 2014), NCBI NR protein database and PSGD. The genes showing significant similarity to the genes in the DroughtDB were listed.

### Diversity analysis in genetic resources

A total of 91 horsegram accessions and one *M. axellare* accession being conserved in CSK-HPAU were used for diversity analysis with ddRAD-Seq reads (Supplementary Table S1). Library construction and variant calling were performed according to Shirasawa et al. (2016). The ddRAD-Seq reads were generated by an Illumina HiSeq 2000 system with reads length of 93 nt, and mapped onto the assembled genome sequences. Calculation of the Jaccard similarity coefficient was conducted using the GGT 2.0 (van Berloo, 2008), and an NJ phylogenetic tree was constructed using MEGA ver 10.1.8 (Kumar et al., 2008). CNVs were identified for HPKM-193 against HPK-4 using CNV-Seq (Xie and Tammi, 2009) with a 1 Mb window.

### Comparative and phylogenetic analysis

The predicted gene sequences were clustered by the CD-HIT program (Fu et al., 2012) with those of *L. japonicus* (Lr_r3.0), *P. vulgaris* (Pvulgaris_218_v1.0), *V. angularis* (Vangularis_v1.a1) and *A. thaliana* (Araport 11). The Ks value was estimated by using KaKs Calculator (Zhang et al., 2006). Macro-synteny between horsegram and *P. vulgaris*, *V. angularis* and *L. japonicus* was investigated based on homologous translated protein sequences searched by BLAST searches with an E-value cut-off of 1E-100. Synteny plots were drawn using the gnuplot program (http://www.gnuplot.info). The phylogenetic analysis was performed by MEGA 7.0.9 beta and TIMETREE based on the single copy genes conserved in horsegram and the four species, *P. vulgaris*, *V, angularis*, *L. japonicus* and *A. thaliana* with 4,152 single copy genes identified by OrthoFinder (Emms and Kelly, 2019). The divergence time of 59 MYA between *L. japonicus* and *P. vulgaris* was used for calibration.

## Data access

The genome assembly data, annotations and gene models are available at the Horsegram Database (http://horsegram.kazusa.or.jp/). The obtained genome sequence reads are available from the DDBJ Sequence Read Archive (DRA) under the BioProject accession number of PRJDB5374.

## Acknowledgement

This work was supported by the Bilateral Joint Research Projects from the Japan Society for the Promotion of Science and Department of Science and Technology of the Government of India, and funds from the Kazusa DNA Research Institute Foundation. We thank A. Watanabe, S. Nakayama, Y. Kishida, M. Kohara, H. Tsuruoka, C. Minami, and S. Sasamoto (KDRI) for their technical assistance. We also thank the National Bureau of Plant Genetic Resources (NBPGR), New Delhi for providing the germplasm lines for diversity analysis.

## Author contributions

The project was designed by S.I., R.C. and T.S. K.S. contributed to the genome sequencing, linkage map and pseudomolecule construction. R.C. contributed to the creation of the mapping population. H.H. contributed to the genome assembly and gene prediction, annotation, and phylogenetic analysis. S.N. contributed to the transcript sequencing. H.N. contributed to the pseudomolecule construction. S.I. contributed to the entire process of data analysis.

## Competing interest

The authors declare no competing interest.

## Notes

http://horsegram.kazusa.or.jp/

### Competing Interest Statement

The authors have declared no competing interest.

