## Supplementary material for "A Chromosome-scale draft genome sequence of horsegram (*Macrotyloma uniflorum*)": Fig. S

**Jellyfish (kmer=17)**

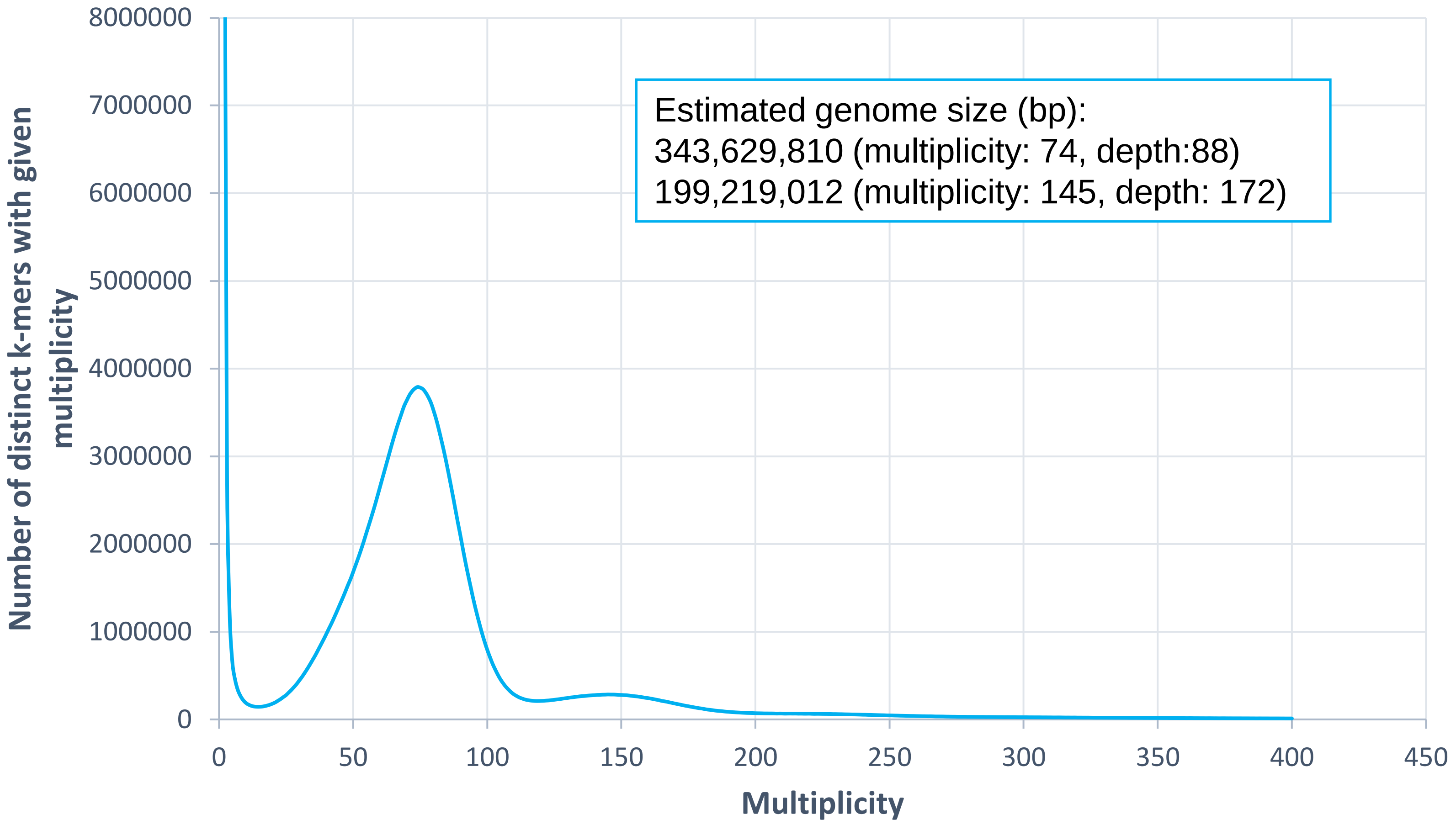


**Fig. S1.** Genome size estimation using Jellyfish with the distribution of the number of distinct kmers (kmer = 17) with the given multiplicity values.

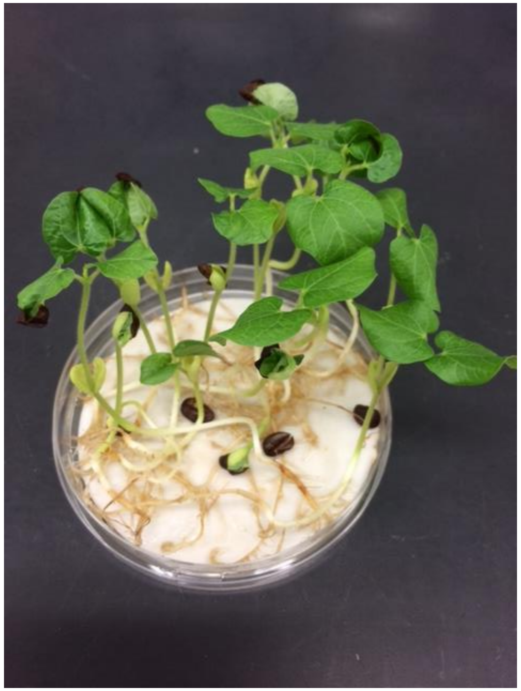
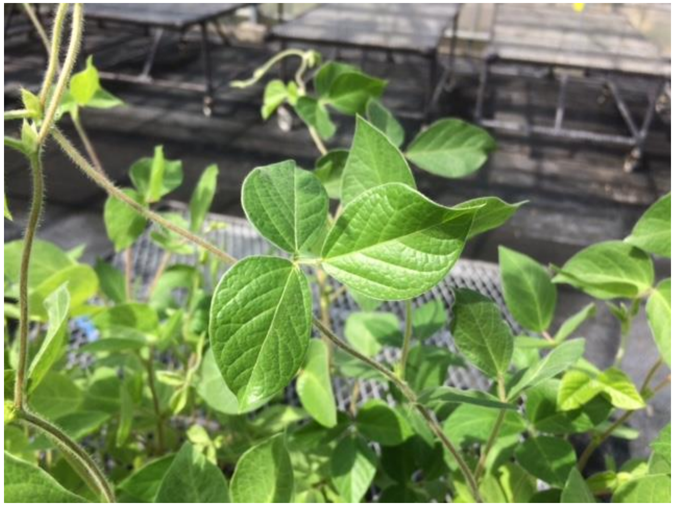
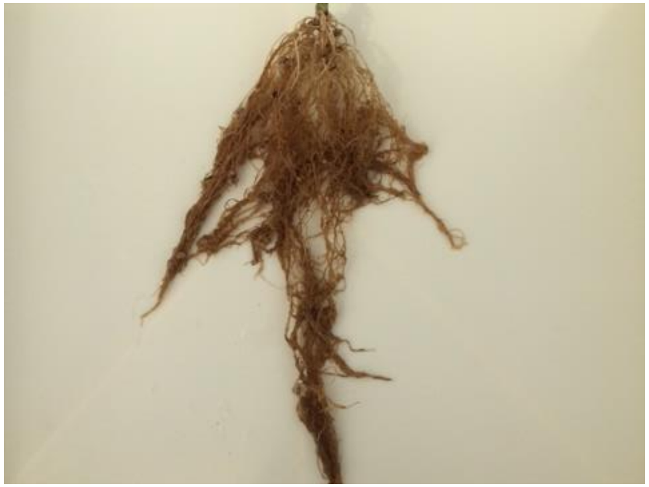
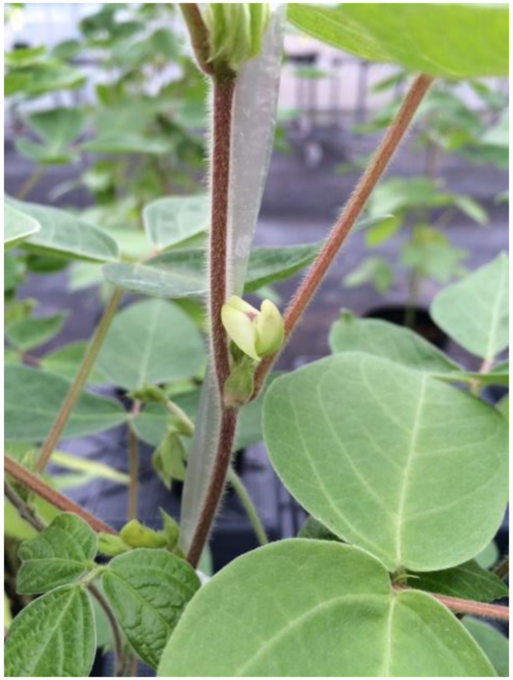
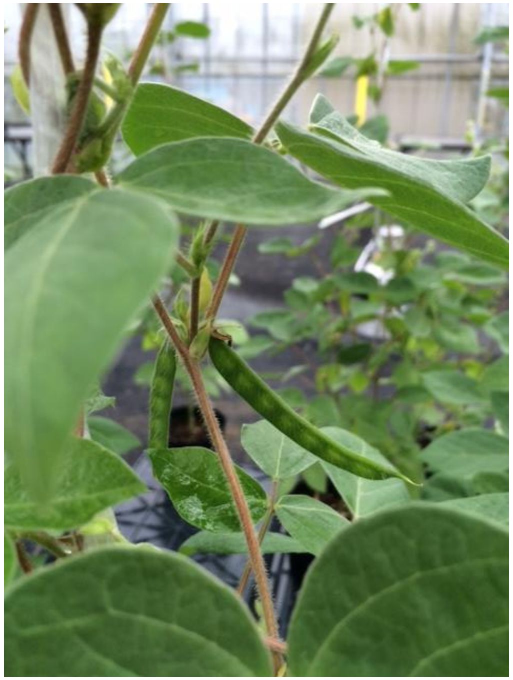

**Fig. S2.** The seedlings, leaves, roots, flowers and young pods of HPK-4 used for Illumina transcript sequencing.

**A:** Numbers and ratios of genes hit against the GO database

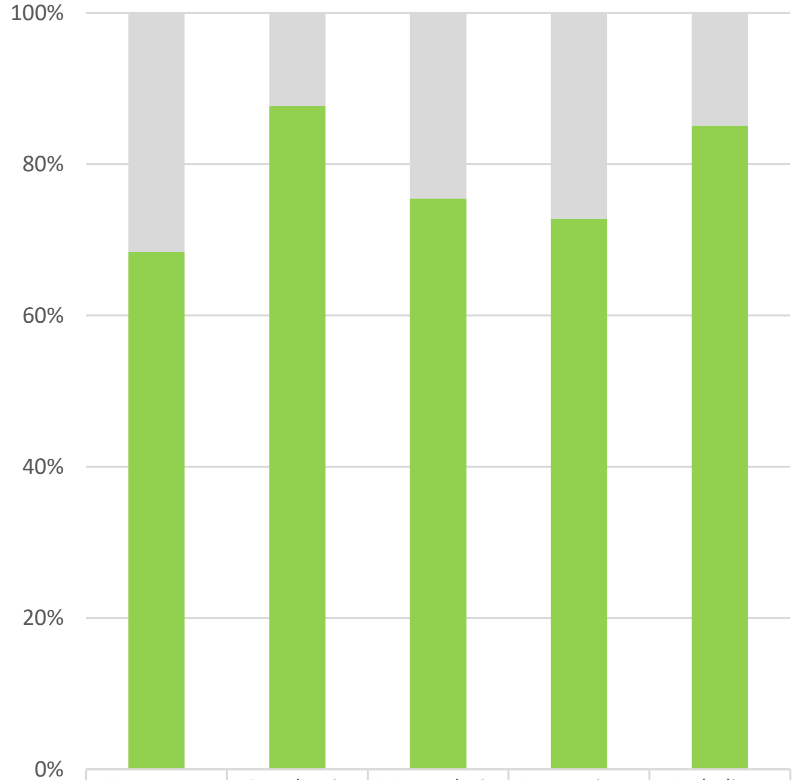


| | Horsegram | P. vulgaris | V. angularis | L. aponicus | A. thaliana |
|---|---|---|---|---|---|
| No hits | 11,406 | 3,348 | 7,661 | 10,817 | 4,123 |
| Hits | 24,699 | 23,849 | 23,580 | 28,917 | 23,532 |

**B:** Ratio of the classified GO functional categories (Root)

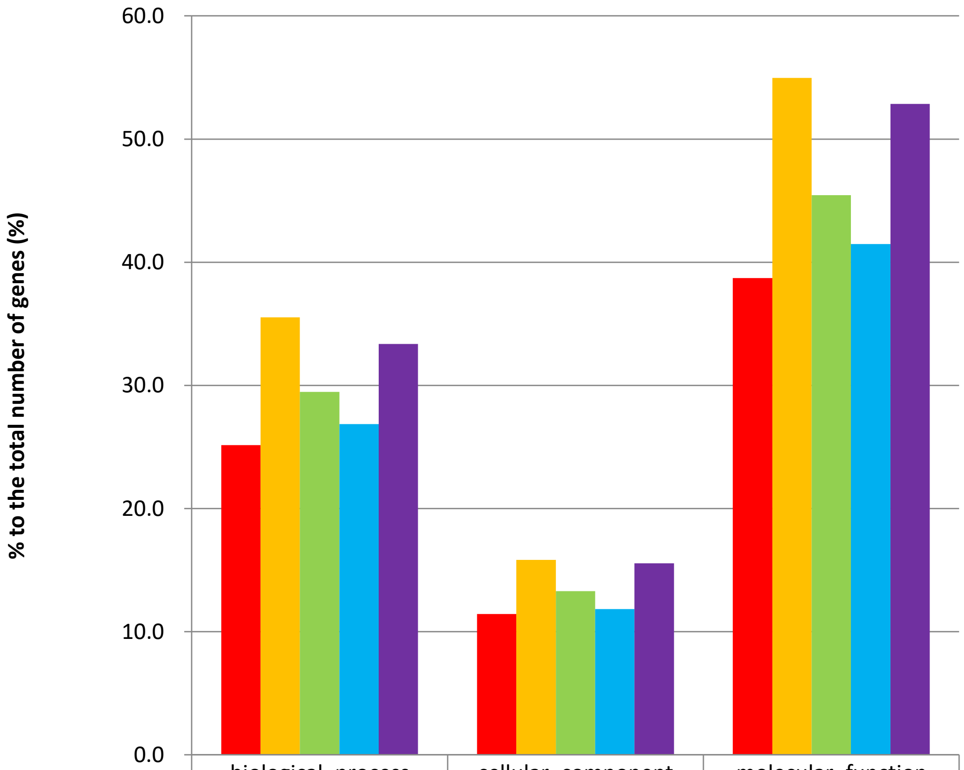


| | biological_process | cellular_component | molecular_function |
|---|---|---|---|
| M. uniflorum | 25.2 | 11.4 | 38.7 |
| P. vulgaris | 35.5 | 15.8 | 55.0 |
| V. angularis | 29.5 | 13.3 | 45.4 |
| L. japonicus | 26.9 | 11.8 | 41.5 |
| A. thaliana | 33.4 | 15.6 | 52.9 |

**Fig. S3. A:** Numbers and ratios of genes annotated by the GO database in horsegram (MUN_r1.1_cds), *P. vulgaris* (Pvulgaris_218_v1.0), *V. angularis* (Vangularis_v1.a1), *L. japonicus* (Lj_r3.0), and *A. thaliana* (Araport11). **B:** Ratios of the classified GO categories in the predicted genes.

**A:** Numbers and ratios of genes hit against the KOG database

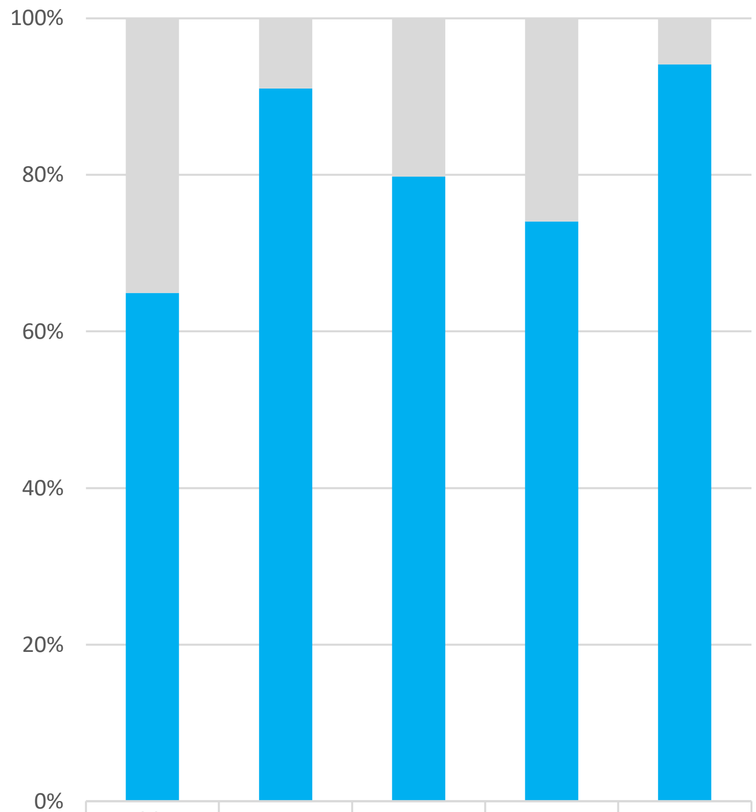


| | M. uniflorum | P. vulgaris | V. angularis | L. japonicus | A. thaliana |
|---|---|---|---|---|---|
| No hits | 10,073 | 2,442 | 6,314 | 10,319 | 1,629 |
| Hits | 18,630 | 24,755 | 24,927 | 29,415 | 26,026 |

**B:** Ratio of the classified KOG functional categories

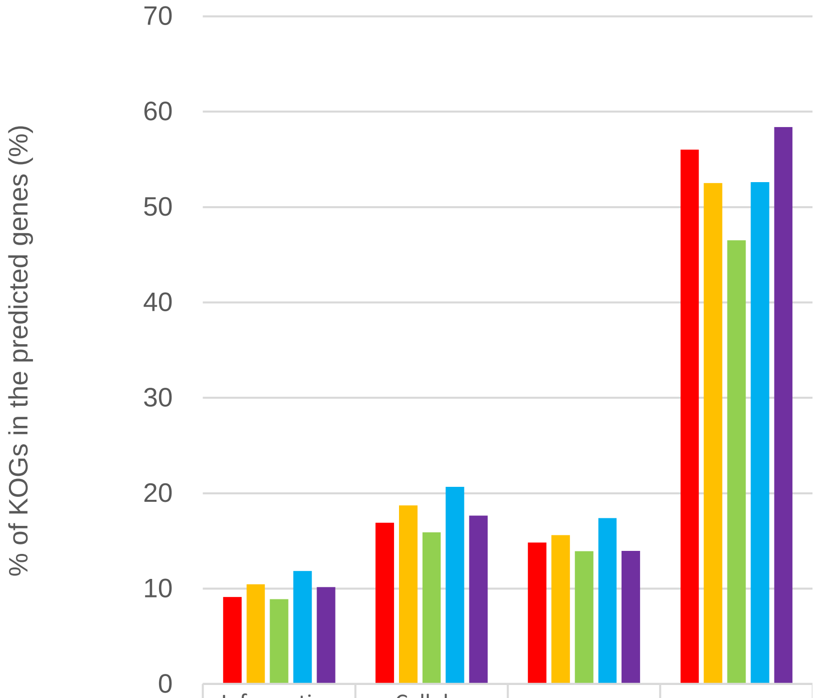


| | Information storage and processing | Cellular processes and signaling | Metabolism | Poorly characterized |
|---|---|---|---|---|
| M. uniflorum | 9 | 17 | 15 | 56 |
| P. vulgaris | 10 | 19 | 16 | 53 |
| V. angularis | 9 | 16 | 14 | 47 |
| L. japonicus | 12 | 21 | 17 | 53 |
| A. thaliana | 10 | 18 | 14 | 58 |

**Fig. S4. A:** Numbers and ratios of genes annotated by the KOG database in horsegram (MUN_r1.1_cds), *P. vulgaris* (Pvulgaris_218_v1.0), *V. angularis* (Vangularis_v1.a1), *L. japonicus* (Lj_r3.0), and *A. thaliana* (Araport11 ). **B:** Ratio of the classified KOG categories in hit genes.

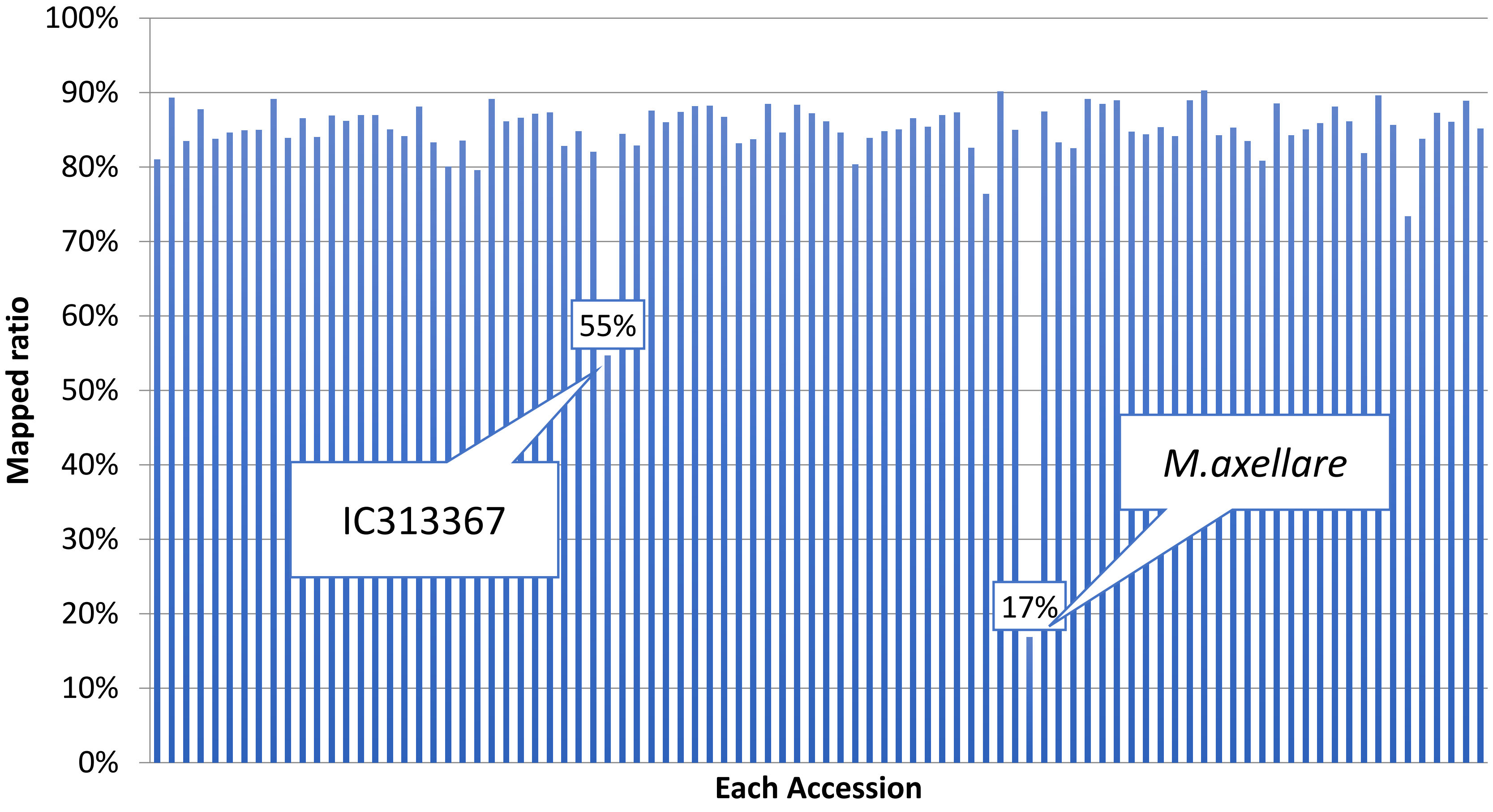


**Fig. S5.** Mapped ratio of the dd-RAD-Seq reads of 92 accessions.
